# Advanced Manufacturing of Coil-Reinforced Multilayer Vascular Grafts to Optimize Biomechanical Performance

**DOI:** 10.1101/2025.01.16.633374

**Authors:** Andrew Robinson, David Jiang, Abbey Nkansah, Juan S. Herrera Duran, Jonathan Leung, Madeline Laude, John Craig, Leopold Guo, Lucas Timmins, Elizabeth Cosgriff-Hernandez

## Abstract

Small diameter vascular grafts require a complex balance of biomechanical properties to achieve target burst pressure, arterial compliance-matching, and kink resistance to prevent failure. Iterative design of our multilayer vascular was previously used to achieve high compliance while retaining the requisite burst pressure and suture retention strength for clinical use. To impart kink resistance, a custom 3D solution printer was used to add a polymeric coil to the electrospun polyurethane graft to support the graft during bending. The addition of this reinforcing coil increased kink resistance but reduced compliance. A matrix of grafts were fabricated and tested to establish key structure-property relationships between coil parameters (spacing, diameter, modulus) and biomechanical properties (compliance, kink radius). A successful graft design was identified with a compliance similar to saphenous vein grafts (4.1 ± 0.4 %/mmHgx10^-2^) while maintaining comparable kink resistance to grafts used currently in the clinic. To explore graft combinations that could increase graft compliance to match arterial values while retaining this kink resistance, we utilized finite element (FE) models of compliance and kink radius that simulated experimental testing. The FE-predicted graft compliance agreed well with experimental values. Although the kink model over-predicted the experimental kink radius values, key trends between graft parameters and kink resistance were reproduced. As an initial proof-of-concept, the validated models were then utilized to parse through a targeted graft design space. Although this initial parameter range tested did not yield a graft that improved upon the previous balance of graft properties, this combination of advanced manufacturing and computational framework paves the way for future model-driven design to further optimize graft performance.

**TOC:** 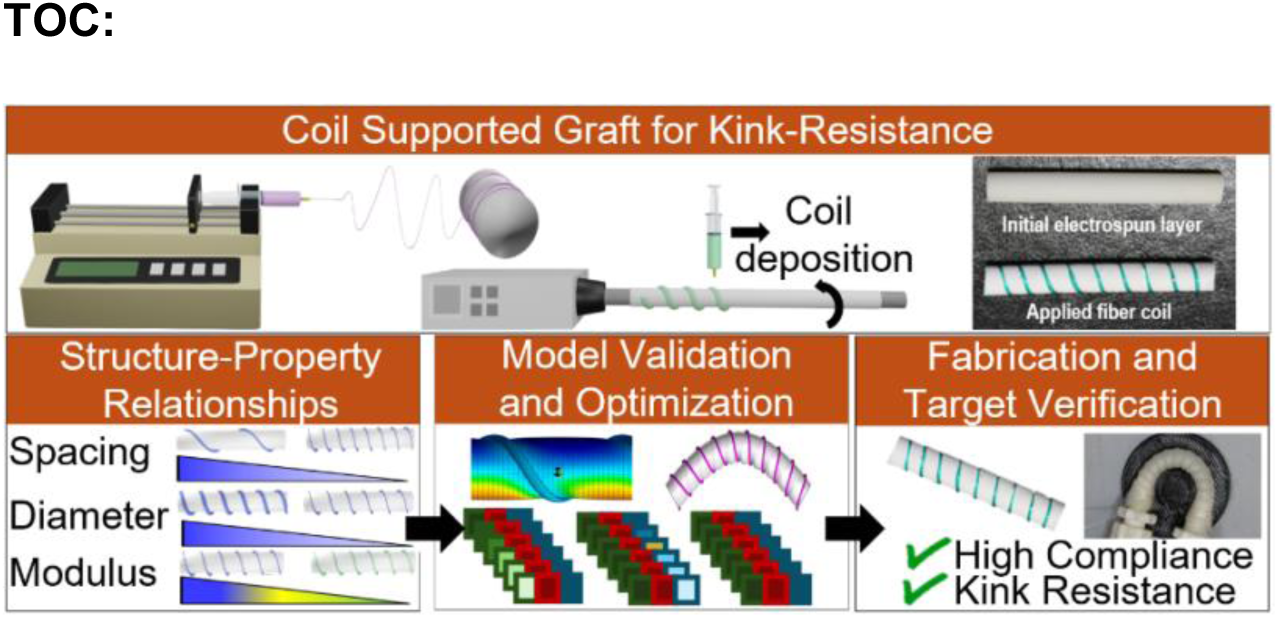

## 1. Introduction

There is an urgent need to develop a synthetic vascular graft for coronary artery bypass grafting (CABG) procedures to address the limited availability of autografts. Multiple grafting procedures, peripheral artery disease, and size mismatch prevent the use of autologous grafts in up to 20% of patients annually.^1^ Although synthetic options such as ePTFE and Dacron have satisfactory outcomes for grafting procedures greater than 6 mm, they have a greater than 40% failure rate within five years in small-diameter procedures due to thrombosis and intimal hyperplasia.^2,3^ Graft thrombosis is attributed to non-specific protein adsorption providing sites for platelet adhesion and activation.^4,5^ Intimal hyperplasia-based failure has been correlated with a mismatch in compliance between the graft and native vessel resulting in hemodynamic alterations and the development of the adverse tissue response.^6,7^

To address these clinical failures, we previously developed a multilayer graft that decouples these disparate biological and mechanical requirements.^1-3,8^ The intimal layer was designed from a bioactive hydrogel with an antifouling substrate based on polyethylene glycol (PEG) for acute thromboresistance.^9,10^ A collagen-mimetic protein derived from group A Streptococcus, Scl2.28 (Scl2)^11^ with integrin-targeting motifs was used to promote endothelialization for sustained thromboresistance.^12,13^ To address intimal hyperplasia, the bioactive hydrogel was reinforced with an electrospun polyurethane mesh to provide requisite burst pressure and improved compliance matching.^3^ This multilayer graft previously demonstrated promising results in both *in vitro* and *ex vivo* models, indicating that high compliance grafts offer potential in overcoming intimal hyperplasia-based failure.^3,13^ Unfortunately, long-term evaluation in a porcine model was prevented due to graft occlusion from kinking. Thus, a method is needed to impart kink-resistance to these multilayer grafts without compromising the burst pressure or high compliance.

External coil supports have been used to clinically utilized to impart kink resistance to synthetic vascular grafts.^14,15^ The coil design can have substantial effects on the resulting graft biomechanical properties.^15-17^ For example, Wu et al. demonstrated that a graft with a tighter coil spacing has improved kink resistance as compared to a wider coil spacing.^15^ Adhikari et al. performed similar studies and demonstrated that a tighter poly(ethylene terephthalate) coil increased the circumferential modulus of the graft.^17^ Although the reinforcing coil can be used to increase kink resistance, the coil also reduces graft compliance due to an increase in circumferential stiffness. To design a graft that balances both high compliance and kink resistance, we must first establish the structure-property relationships between coil parameters (e.g., spacing, diameter, modulus) and biomechanical properties. However, the design space between the different coil parameters is exceedingly large to explore iteratively. Thus, a method is needed to rapidly parse the expansive design space to identify promising graft designs and inform fabrication directions.

Computational simulations have been utilized to predict device performance across the cardiovascular space.^18-20^ For example, Tamimi et al. developed computational simulations to predict the thickness, crosslinking time, and ratios of polycaprolactone (PCL)/gelatin and PCL/tropoelastn required to achieve target compliances for a tissue engineered vascular graft.^18^ Cosgriff-Hernandez and Timmins highlighted the potential of model directed fabrication to accelerate development by noting that experimentally evaluating an 80 parameter design space would take 6-9 months compared to *in silico* evaluations that took ∼4 hours.^21^ We previously developed compliance and kink models that simulate the experimental compliance and kink radius testing methods to evaluate the effect of coil design on these two competing mechanical properties.^22^ This enables batch processing of the design space for rapid identification of optimized graft designs *in silico* that can be fabricated using our advanced manufacturing methods.

In this work, we utilized advanced manufacturing to fabricate a coil-reinforced multilayer vascular graft with improved compliance and kink resistance. Reinforcing coils were added to the electrospun polyurethane graft using a custom 3D solution deposition adapted from Li et al.^16^ A series of experiments were performed to establish structure-property relationships between coil parameters (spacing, diameter, modulus) and graft biomechanical properties (compliance, kink resistance). Experimental values from graft testing were then used to validate finite element (FE) models to predict graft compliance and kink resistance. These validated models were used to parse a targeted design space *in silico* to identify candidate graft designs. Collectively, this work utilized coil-reinforcement to design a kink-resistant multilayer graft and explore a computational framework to test graft design parameters *in silico* for graft optimization.

## 2.0 Materials and Methods

All reagents were obtained and utilized as received from VWR (Radnor, PA) or Sigma-Aldrich (Milwaukee, WI) unless stated otherwise.

### 2.1 Electrospun Graft Fabrication

Control grafts were fabricated via electrospinning utilizing a 14 wt% solution of Bionate® 80A (DSM Biomedical) in a solvent mixture of 70:30 dimethylacetamide:tetrahydrofuran. The solution was passed at a rate of 0.5 ml/hr through a 20-gauge blunt needle that was charged to +18.4-20.5 kV. A 5 mm mandrel was dip-coated in a solution of 5 wt% polyethylene glycol (PEG; 35 kDa) in dichloromethane to facilitate graft removal following spinning. The collecting mandrel was placed 50 cm from the needle, charged to -2.5 kV, and rotated at 500 RPM. Relative humidity and temperature were 40-47% and 23.5-25.4 °C, respectively. The graft was allowed to dry overnight and then removed by placing the mandrel in water for ∼30 minutes to dissolve the PEG layer and sliding the graft off. Grafts were trimmed to ∼4 cm and mesh thickness (n = 4 per graft, grafts = 3) was measured using force-normalized calipers, Mitutoyo 547-500S.

### 2.2 Coil-Supported Graft Fabrication

Coil-supported grafts were fabricated in three layers using a combination of electrospinning and a custom-developed 3D printer, **Figure 1A**. The first electrospun layer was fabricated from an 18wt% solution of Bionate® 80A in a solvent mixture of 70:30 dimethylacetamide:tetrahydrofuran. The solution was passed through a 20-gauge blunt needle at 0.5 ml/hr charged to +16.5-22.5 kV. Fibers were collected on a 5 mm PEG-coated mandrel, as described above. The mandrel was rotated at 500 RPM and charged to –2.5 kV with a working distance of 50 cm. Temperature and relative humidity during the spin were 23.2-25.1 °C and 42-48%, respectively. An initial mesh thickness of ∼120-150 µm was targeted for the first layer. A polymeric coil was then applied to the grafts via solution deposition with a custom coil printer to control coil spacing, diameter, and modulus, **Figure S1 and Table S1**. Coil spacing was controlled by altering the lateral speed of the printer. Coil diameter was controlled through a combination of needle gauge and flow rate of the polymeric solution. Coil modulus was altered by using either Bionate® 55D (18wt% in 1,1,1,3,3,3-Hexafluoro-2-propanol, SynQuest Laboratories) with a modulus of 8.1 MPa or Bionate® 80A (21.5wt% in 1,1,1,3,3,3-Hexafluoro-2-propanol, SynQuest Laboratories) with a modulus of 2.9 MPa.^22^ The coiled graft was left to dry at ambient conditions in a fume hood. Following drying, a second electrospun layer was deposited using the 18wt% Bionate® 80A solution. The solution was passed through a 20-gauge needle at a flow rate of 0.5ml/hour with +15.3-22.1 kV applied to the needle tip. A working distance of 50 cm was used with a -2.5 kV applied to the mandrel. The temperature and humidity were 23.4-26.1 °C and 41-47%, respectively. Grafts were allowed to air dry on the mandrel overnight and removed as described prior. Mesh thickness (n = 4 per graft, grafts = 3) and coil dimensions (n = 5 per graft, grafts 3) were measured using force-normalized calipers, Mitutoyo 547-500S, with the minor diameter of the coil being refered to as the coil diameter hence forth. Coil spacing was measured in ImageJ from the middle to of the coil to the middle of the adjacent coil (n = 5 per graft, grafts = 3)

**Figure 1:**
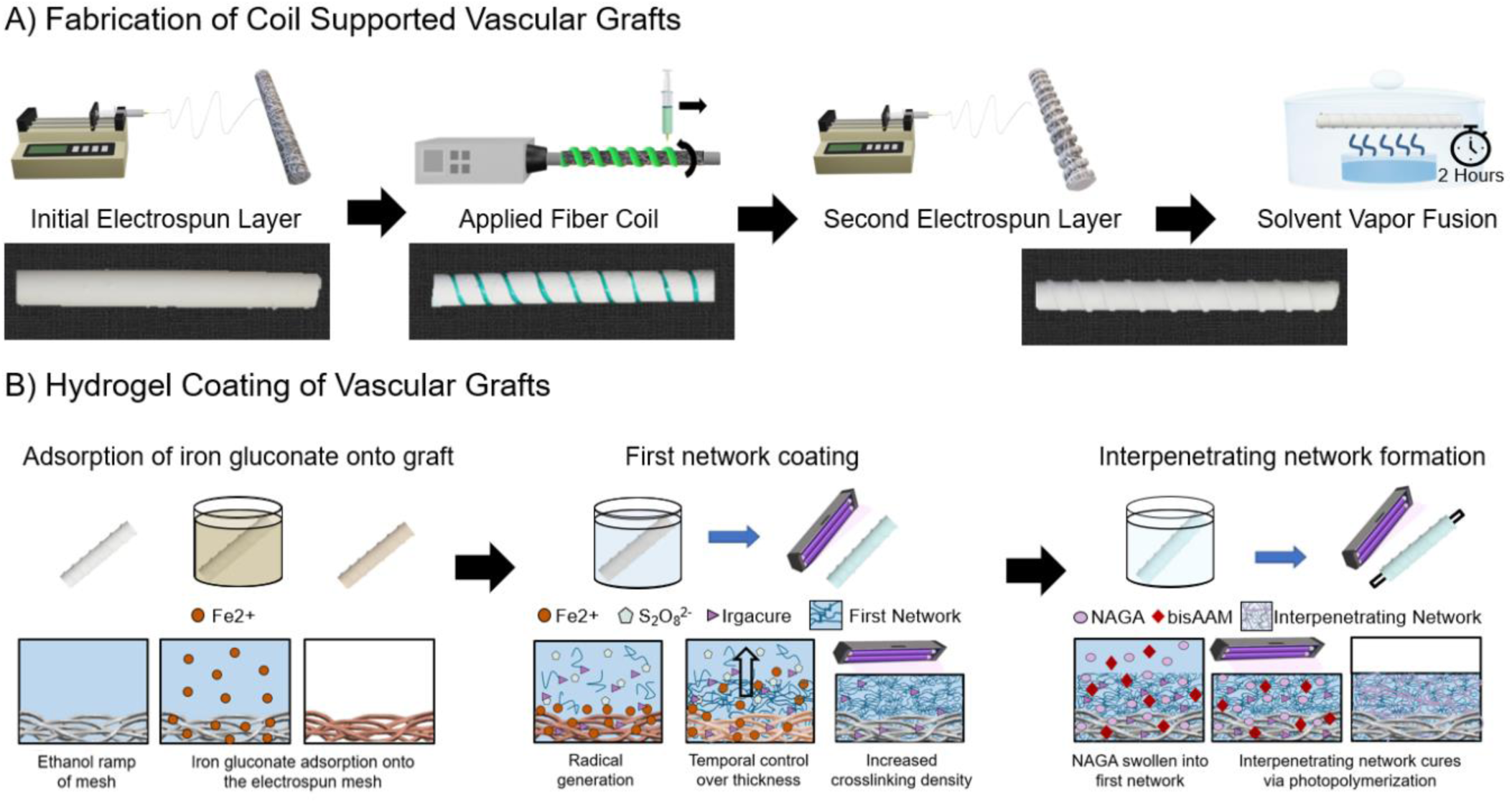
Fabrication of the coil-supported vascular graft with hydrogel coating. A) Fabrication of the electrospun sleeve via an initial electrospun layer, followed by coil deposition using a custom solution printer, and a final electrospun layer. Solvent vapor welding is performed for two hours using tetrahydrofuran to improve construct integrity. B) Hydrogel coating of electrospun grafts performed with diffusion-mediated redox initiated crosslinking of the PEUDAm first network that sets the thickness of the hydrogel coating layer. NAGA, bisAAm, and photoinitiators are then swollen into the first network and cured via photoinitiation to form the final interpenetrating network hydrogel coating.

### 2.3 Solvent Vapor Welding and Graft Characterization

To improve integrity between the electrospun and coil layers, both electrospun control grafts and coil-supported grafts underwent solvent vapor welding. Following removal of the grafts from the electrospinning mandrel, a 5 mm Teflon rod was placed inside the grafts and the grafts were allowed to air dry overnight. Grafts then underwent solvent vapor exposure for two hours, **Figure 1A**. A 12 cm diameter crystallization dish with 40 ml of tetrahydrofuran was placed inside a 17 cm diameter desiccator. The grafts were placed on a grate over the crystallization dish and the desiccator was closed. Following the two-hour solvent vapor fusion, the grafts were dried under vacuum overnight. Fiber diameter was anaylzed via scanning electron microscopy (Phenom Pro, NanoScience Instruments, Phoenix, AZ,). Samples were coated with 5 nm of gold (Sputter Coater 108, Cressington Scientific Instruments, Hertfordshire, UK) and images at 5000X magnification. Average fiber diameter was characterized in ImageJ by drawing a mid-line and measuring the diameter of the first ten fibers that crossed the line (n = 5 images per graft, graft = 3, n = 150 per graft).

### 2.4 Hydrogel Coating

Synthesis of polyether urethane diacrylamide (PEUDAm) 20 kDa was performed using established protocols.^10^ The hydrogel coating of electrospun polyurethane grafts was accomplished through diffusion-mediated redox initiated crosslinking, following established protocols, **Figure 1B**.^11^ Electrospun grafts were treated with a graded ethanol/water wetting ladder (70 vol%, 50 vol%, 30 vol%, and 0 vol% ethanol in water, with each concentration applied for 15 minutes). Following wetting, grafts were submerged in an iron gluconate dihydrate solution (IG, 3 wt/vol% [Fe^2+^], as determined with the Ferrozine Assay^23^) for 15 minutes. After a brief dip in methanol, the substrates were air-dried for one minute then immersed in aqueous solutions of 10 wt/vol% PEUDAm 20 kDa with ammonium persulfate (APS, 0.05 wt/vol%) and 2 wt/vol% Irgacure 2959 for 20 seconds. Following dip coating, grafts were placed on a UV plate (Intelligent Ray Shuttered UV Flood Light, Integrated Dispensing Solutions, Inc., 365 nm, 4 mW/cm^2^) for 12 minutes for additional photoinitiated crosslinking of the PEUDAm hydrogel layer. Coated grafts were then dried overnight and soaked for 5 hours in DI water with water exchanges performed after 10 minutes, 1 hour, and 3 hours to remove unreacted reagents. Grafts were dried completely prior to soaking in a solution of 20 wt/vol% N-acryloyl glycinamide (NAGA, BLD Pharma), 0.1 mol% bisacrylamide (bisAAm), and 2 wt/vol% Irgacure 2959 overnight protected from light at 4 °C for incorporation of the second network prepolymers. Grafts were then placed on a 4 mm glass rod to prevent bending during curing and mounted to a rotating device. The rotating device was then set inside a custom-made mirror apparatus placed on a UV transilluminator to promote an even UV cure of the final interpenetrating network (IPN) hydrogel. Composites were air-dried overnight and soaked in DI water to remove sol fraction.

### 2.5 Vascular Graft Mechanical Testing

Compliance was performed as previously described.^3^ Briefly, the ends of the composite grafts were fixed to a barb fitting with a monofilament line and pressurized from 0 mmHg to 120 mmHg at a rate of 2 ml/min using a syringe pump. The diameter of the grafts was measured at 80 mmHg and 120 mmHg using a laser micrometer (Z-Mike 1210 Laser Micrometer). Compliance was calculated according to **Eq. 1** where D_80_ is the lumen diameter of the graft at 80 mmHg, D_120_ is the lumen diameter of the graft at 120 mmHg, and ΔP is the pressure change (40 mmHg). Compliance was measured at the locations of greatest compliance between the coils (n = 8 per graft, grafts = 3). Effect of compliance and location are included in **Figure S2**.

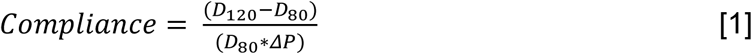

Kink radius tests were performed using 3D printed mandrels that were developed in accordance with ISO 7198:2016 A5.8.^15,24^ Composite control and coil-supported grafts were secured to barb fittings and pressurized to 100 mmHg at a flow rate of 2ml/min. The grafts were bent to the curvature of each mandrel and imaged using a DSLR camera. The diameter loss was measured in ImageJ according to **Eq. 2** where D_o_ is the patent diameter of the unbent graft that had been pressurized to 100 mmHg and D_m_ is the patent diameter of the graft during bending at a specific mandrel. Kink radius was defined as the radius of the mandrel at a which there was 50% or greater loss in diameter of the graft (n = 3 per grafts, grafts = 3).^15^

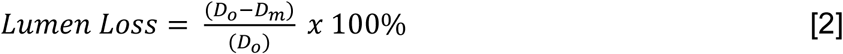

Burst pressure and suture retention strength were assessed as previously described.^3^ Briefly, a latex liner was inserted into the grafts that were then fixed to barb fittings to assess burst pressure. Grafts were pressurized at a flow rate of 50 ml/min with the flow obstructed. The highest pressure measured was recorded as the burst pressure of the grafts (n = 3). For suture retention strength, grafts were cut lengthwise, and a suture was looped through the mesh 2 mm from the edge of the graft perpendicular to the direction of tension. The graft end was trimmed to remove the coil from the area that suture would pull through prior to the application of the suture. The graft and suture were then secured to pneumatic clamps. A uniaxial strain rate of 100 mm/min was applied using an Instron 3345 with the maximum observed force recorded as the suture retention strength (n = 2 per graft, grafts = 3).

### 2.6 Model Validation and Optimization for Compliance and Kink Resistance

FE models of the graft were developed using the open-source FEBio software.^25,26^ The framework to automatically create and discretize the model geometries, apply model definitions (i.e., boundary conditions, loads, contact), and generate FEBio input files are detailed in Jiang et al.^27^ Briefly, the automated scheme utilized the open-source GIBBON toolbox to construct and discretize the graft and coil geometries.^28^ The graft and coil material properties were derived from mechanical testing and Bionate® PCU product sheets (DSM Biomedical Inc) and modeled in FEBio as an uncoupled neo-Hookean hyperelastic material with nearly incompressible behavior. Contact interfaces were defined to connect the coil to the graft outer-surface and to prevent self-penetration of the elastic bodies during deformation. To predict compliance (C_graft_), both graft ends were fixed in all directions to prevent rigid body motion. Next, the graft was pressurized to 80 and 120 mmHg. Nodal values for position were extracted, and compliance was calculated according to **Eq. 1**. To predict kink resistance (K_graft_), the graft model was lengthened to 8 cm to reduce the edge effects. Both graft ends were fixed in all directions and subsequently pressurized to 100 mmHg. Next, both ends were equally rotated by 90° in opposite directions and displaced downwards and inwards to generate an effective bending moment, as previously performed.^29^ Nodal values for position were extracted, and the kink radius was calculated according to **Eq. 2**.

### 2.8 Statistical Analysis

Data are displayed as mean ± standard deviation. Statistical analysis was performed using either a t-test when two groups were compared, or an analysis of variation (ANOVA) with a Tukey’s multiple comparison test when three were compared. Statistical significance was set at p < 0.05.

## 3.0 Results

### 3.1 Fabrication

Coil-supported vascular grafts were fabricated using a combination of electrospinning and 3D solution deposition, **Figure 2**. Fiber diameter of the electrospun grafts was 1.9 ± 0.4 µm. Solution deposition of the coil resulted in a flattening of the coil from circular to ovular with the width (major diameter) being 2.19 ± 0.09 times the height (minor diameter). The thickness of the electrospun mesh and hydrogel were 0.26 ± 0.01 mm and 0.15 ± 0.02 mm, respectively.

**Figure 2:**
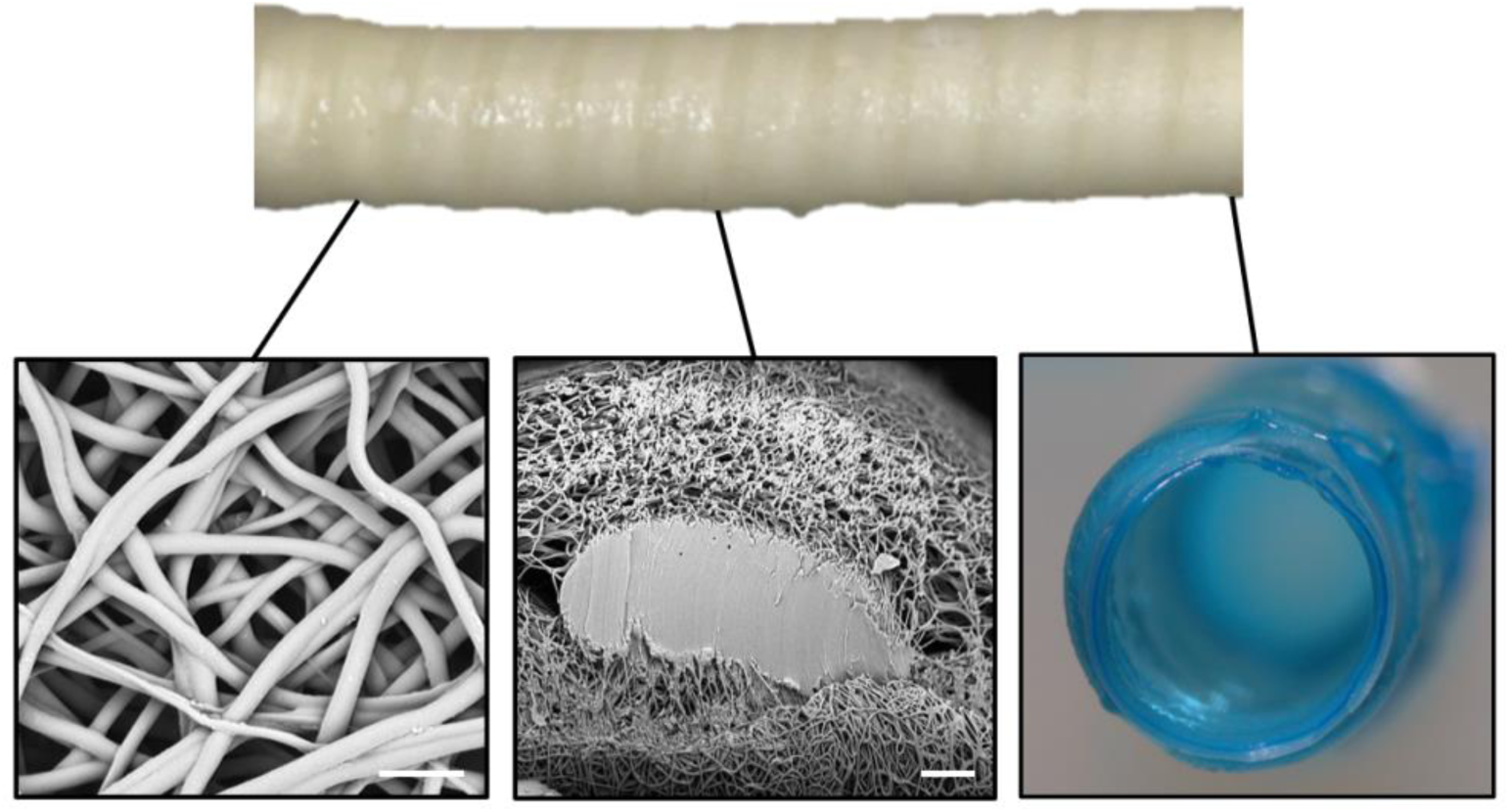
Composite graft with coil reinforcement. Scanning electron micrograph of the electrospun fibers (scale bar equal 10 µm). Cross-section of the electrospun sleeve and embedded coil support (scale bar is 50 µm). Cross-section of a composite graph depicting the electrospun mesh and hydrogel coating.

### 3.2 Effect of Coil Spacing on Compliance and Kink Radius

Coil spacing was investigated to determine the effect on graft compliance and kink radius, **Figure 3**. A coil of Bionate® 55D (modulus = 8.1 MPa) was used with target coil spacings of 2.5 mm, 5.0 mm, and 7.5 mm. Coil spacing was altered by changing the lateral speed of the custom 3D printer and fabricated graft coil spacing (2.5 ± 0.1 m, 5.1 ± 0.2 mm, 7.6 ± 0.2 mm) were in good agreement with target values. Coil diameter was maintained across all graft designs with 0.34 ± 0.01 mm, 0.33 ± 0.01 mm, 0.34 ± 0.01 mm, respectively. Graft compliance was shown to decrease with tighter coil spacing, **Figure 3B**. For example, the grafts with a coil spacing of 7.5 mm had a compliance of 6.3 ± 0.8 %/mmHg x 10^-2^ in comparison to the grafts with a spacing of 2.5 mm that had a compliance of 3.4 ± 0.5 %/mmHg x 10^-2^. This represents a 46% decrease in compliance. A decrease in coil spacing also resulted in a corollary decrease in kink radius, **Figure 3C**. Grafts with a 7.5 mm coil spacing had a kink radius of 11.8 ± 1.5 mm; whereas, grafts with a coil spacing of 2.5 mm had a kink radius of 3.8 ± 0.5 mm. This represents a 68% decrease in kink radius.

**Figure 3:**
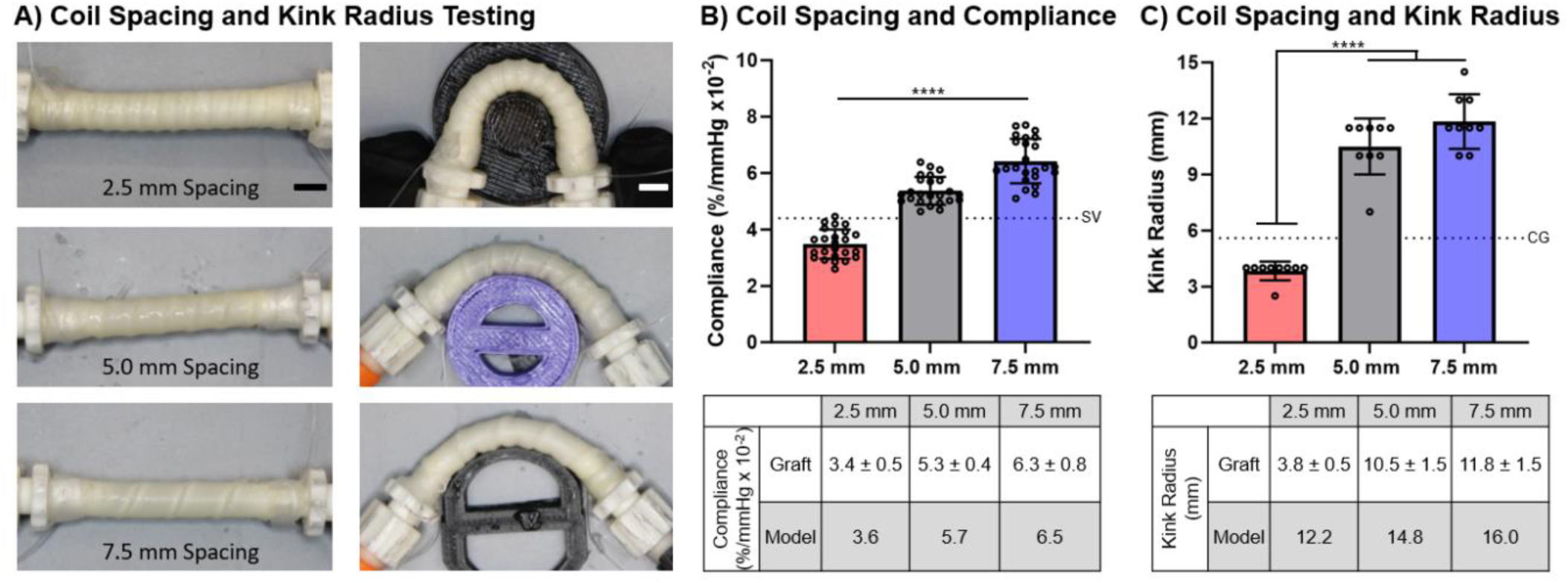
Effect of coil spacing (2.5, 5.0, and 7.5 mm) on graft compliance and kink radius with coil diameter (0.35 mm) and modulus (8.1 MPa) held constant. A) Representative images of coil-supported graft prior to and during the kink radius test. Representative kink images are of the template prior to kinking. Scale bar is 5 mm. B) Effect of coil spacing on graft compliance with comparison to model predictions. SV denotes the compliance of the saphenous vein.^30^ C) Effect of coil spacing on graft kink radius with comparison to model predictions. CG denotes the kink radius of clinical synthetic grafts.^31^ **** denotes statistical significance between all groups using an ANOVA with a Tukey’s multiple comparisons test (p < 0.0001).

### 3.3 Effect of Coil Diameter on Compliance and Kink Radius

To investigate the effect of coil diameter on graft biomechanical properties, coil diameter was varied (0.15, 0.25, 0.35 mm) while holding coil spacing (2.5 mm) and modulus constant (8.1 MPa), **Figure 4**. Coil diameter was controlled by a combination of needle gauge and flow rate of the extruded coiling solution. Fabricated coil diameter and spacing matched target diameters with 0.15 ± 0.01 mm, 0.25 ± 0.01 mm, and 0.34 ± 0.01 mm with coil spacings maintained at 2.6 ± 0.1 mm, 2.6 ± 0.1 mm, and 2.5 ± 0.1 mm, respectively. Increasing coil diameter resulted in a reduction in graft compliance. A coil diameter of 0.15 mm resulted in a graft compliance of 4.9 ± 0.5 %/mmHgx10^-2^ as compared to a coil diameter of 0.35 mm that reduced graft compliance to ± 0.5 %/mmHgx10^-2^. This represents a 30% decrease in compliance. Increasing coil diameter also reduced graft kink radius. A coil diameter of 0.15 mm resulted in a graft kink radius of 8.8 ± 0.7 mm as compared to a coil diameter of 0.35 mm that reduced graft kink radius to 3.8 ± 0.5 mm. This represents a 57% decrease in kink radius.

**Figure 4:**
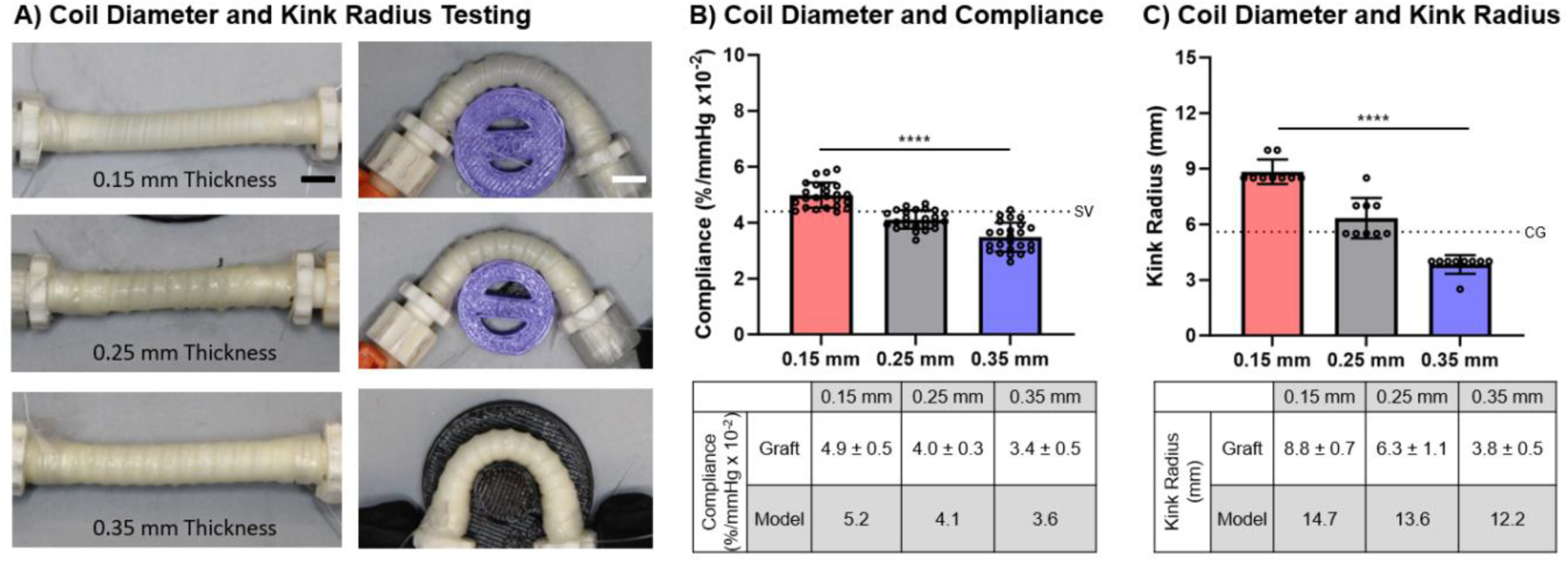
Effect of coil diameter (0.15, 0.25, and 0.35 mm) on compliance and kink radius with spacing (2.5 mm) and modulus (8.1 MPa) held constant. A) Representative images of coil-supported grafts prior to and during the kink radius test. Representative kink images are of the template prior to kinking. Scale bar equals 5 mm. B) Effect of coil diameter on graft compliance with comparison to model predictions. SV denotes the compliance of the saphenous vein.^30^ C) Effect of coil diameter on graft kink radius with comparison to model predictions. CG denotes the kink radius of clinical grafts.^31^ **** denotes statistical significance between all groups using an ANOVA with a Tukey’s multiple comparisons test (p < 0.0001).

### 3.4 Effect of Coil Modulus on Compliance and Kink Radius

To investigate the effect of coil modulus on graft biomechanical properties, coil modulus was varied while holding coil spacing (2.5 mm) and diameter constant (0.35 mm), **Figure 5**. Coil modulus was varied by using either Bionate® 55D (modulus = 8.1 MPa) or Bionate® 80A (2.9 MPa) for the coiling polymer. Coil diameter and spacing were 0.33 ± 0.01 mm and 2.6 ± 0.1 mm for Bionate® 55D and 0.35 ± 0.01 mm and 2.6 ± 0.1 mm for Bionate® 80A. Increasing coil modulus resulted in a reduction of compliance from 4.1 ± 0.4 %/mmHgx10^-2^ to 3.4 ± 0.5 %/mmHgx10^-2^ and a small decrease in kink radius from 4.8 ± 1.5 mm to 3.8 ± 0.5 mm. This represents a 17% decrease in compliance and a 21% decrease in kink radius.

**Figure 5:**
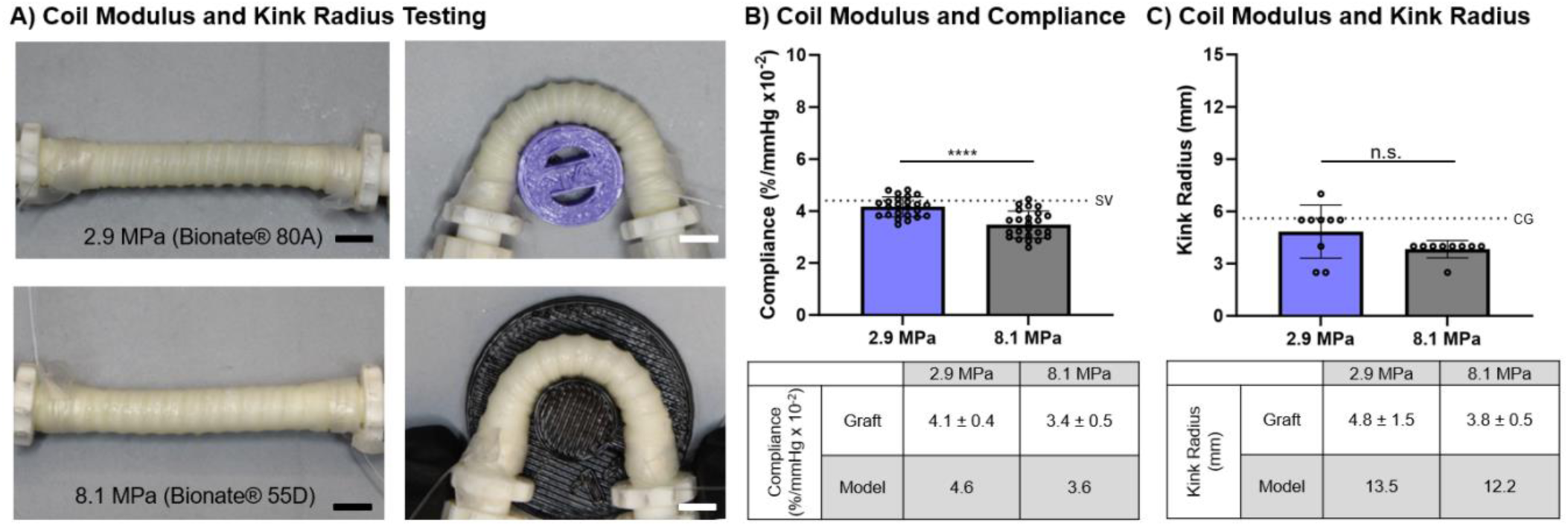
Effect of coil modulus (2.9 MPa, 8.1 MPa) on compliance and kink radius with spacing (2.5 mm) and diameter (0.35 mm) held constant. A) Representative images of coil-supported graft prior to and during the kink radius test. Representative kink images are of the template prior to kinking. Scale bar equals 5 mm. B) Effect of coil modulus on graft compliance with comparison to model predictions. SV denotes the compliance of the saphenous vein.^30^ C) Effect of coil modulus on graft kink radius with comparison to model predictions. CG denotes the kink radius of clinical grafts.^31^ **** denotes statistical significance using a t-test (p < 0.0001). N.S. indicates no significance using a t-test (p > 0.05).

### 3.5 Biomechanical Properties of Coil-reinforced Multilayer Graft

Structure-property relationships were used to identify a coil-reinforced multilayer graft that met target design parameters for potential clinical use. Namely, minimum compliance, burst pressure, and suture retention strengths similar to the saphenous vein and a kink radius in the range of commercial grafts, **Figure 6**. The graft had a coil-support with a modulus of 2.9 MPa, a spacing of 2.5 mm, and a diameter of 0.35 mm. The coil-reinforced graft displayed a marked improvement in kink resistance as indicated by a reduction in kink radius to 4.8 ± 1.5 mm from 11.7 ± 1.4 mm of the control graft. However, there was also a loss in graft compliance from 8.1 ± 0.5 %/mmHg x 10^-2^ for the control graft to 4.1 ± 0.4 %/mmHg x 10^-2^ due to the coil support. This kink radius is in the range of synthetic grafts in clinical use (≤ 5.6 mm) and the graft compliance is comparable to the saphenous vein (∼4.4 %/mmHg x 10^-2^).^31,32^ Burst pressure and suture retention strength of the candidate graft were investigated to ensure it met surgical requirements. The coil-reinforcement increased graft burst pressure to 1767 ± 145 mmHg as compared to 1219 ± 181 mmHg of the control graft. There was no significant difference in suture retention strength of the coil-reinforced graft (249 ± 51 gF) and the control graft (256 ± 13 gF). Both grafts exceeded reported burst pressure and suture retention strength reported for the saphenous vein, 984 mmHg and 203 gF, respectively.^3,33,34^

**Figure 6:**
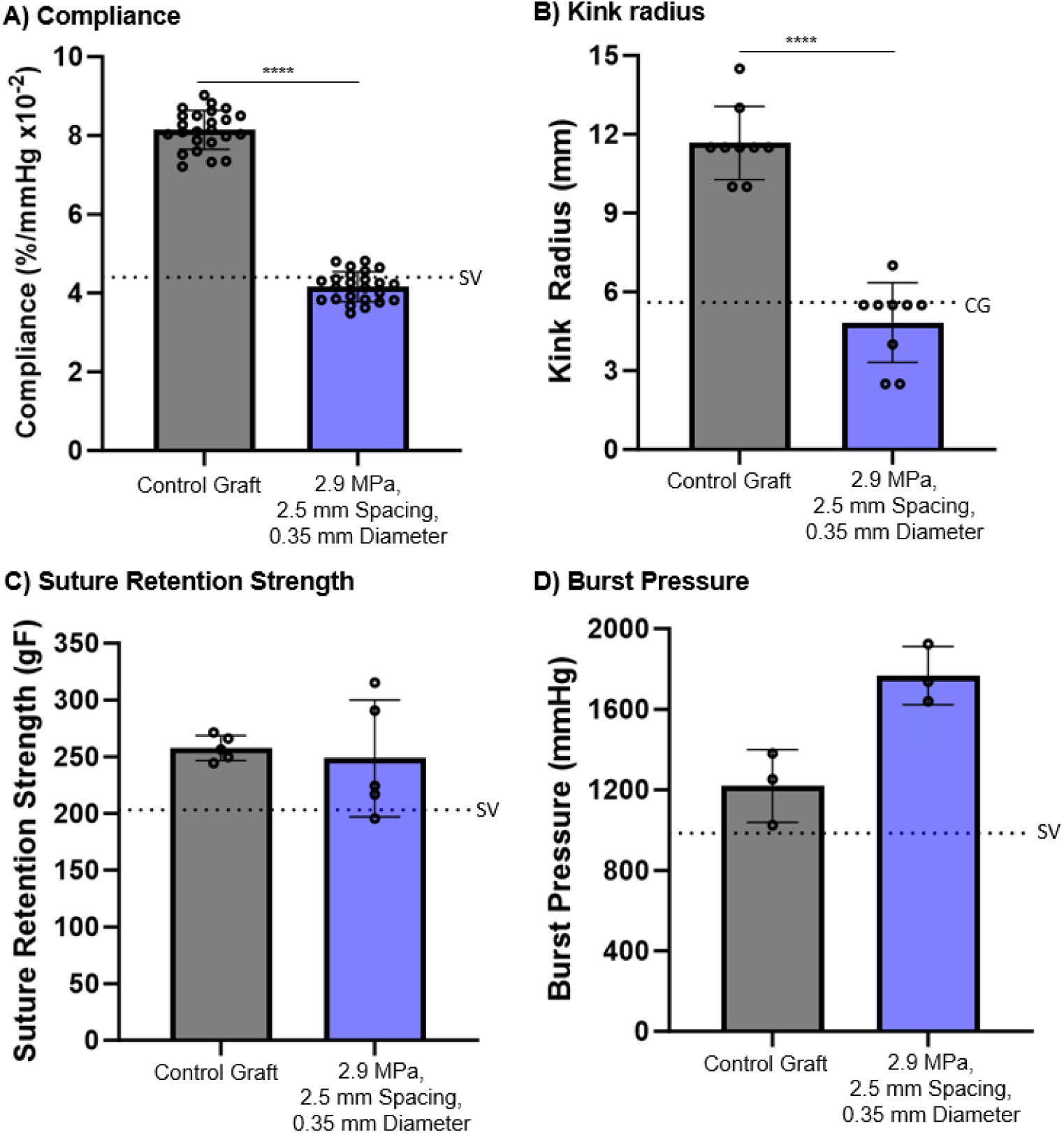
Biomechanical properties of the control and candidate coil-reinforced graft (coil modulus = 2.9 MPa, spacing = 2.5 mm, diameter = 0.35 mm). A) Compliance, SV denotes the compliance of the saphenous vein;^30^ B) Kink radius, CG denotes the kink radius of clinical grafts;^31^ C) Suture retention strength, SV denotes the suture retention strength of the saphenous vein;^30^ D) Burst pressure, SV denotes the burst pressure of the saphenous vein.^30^ **** denotes statistical significance using a t-test (p < 0.0001).

### 3.4 Computational Framework for *In Silico* Testing

Graft compliance and kink radius FE models were previously developed to predict graft biomechanical properties as a function of coil design, **Figure 7**.^27^ The graft compliance model demonstrates how the coil restricts graft expansion along areas of coil contact during pressurization at pressures of 0 mmHg, 80 mmHg, and 120 mmHg. The graft kink radius model bends the graft along a pre-determined path to mimic standard experimental testing. The reduction in the diameter of the graft was measured and the time point of a kink and subsequent kink radius were determined as described previously.^27^ The simulation results were compared to experimental results for validation of the model predictions using a linear regression. The max difference between graft compliance FE model predictions was less within 11% of experimental results. The linear regression of the graft compliance model had a slope of 0.95 and a r^2^ value of 0.99 demonstrating high predictive capability, **Figure 7B**. The graft kink radius model overpredicted the kink radius as compared to experimental results. For example, a graft with a coil modulus of 8.1 MPa, a diameter of 0.35 mm, and a spacing of 2.5 mm had a kink radius of 3.8 mm experimentally; whereas, the model predicted a graft kink radius of 12.2 mm. Although the model overpredicted the graft kink radius, it accurately predicted trends as noted by the linear regression with a slope of 0.40 and an r^2^ value of 0.92, **Figure 7D**.

**Figure 7:**
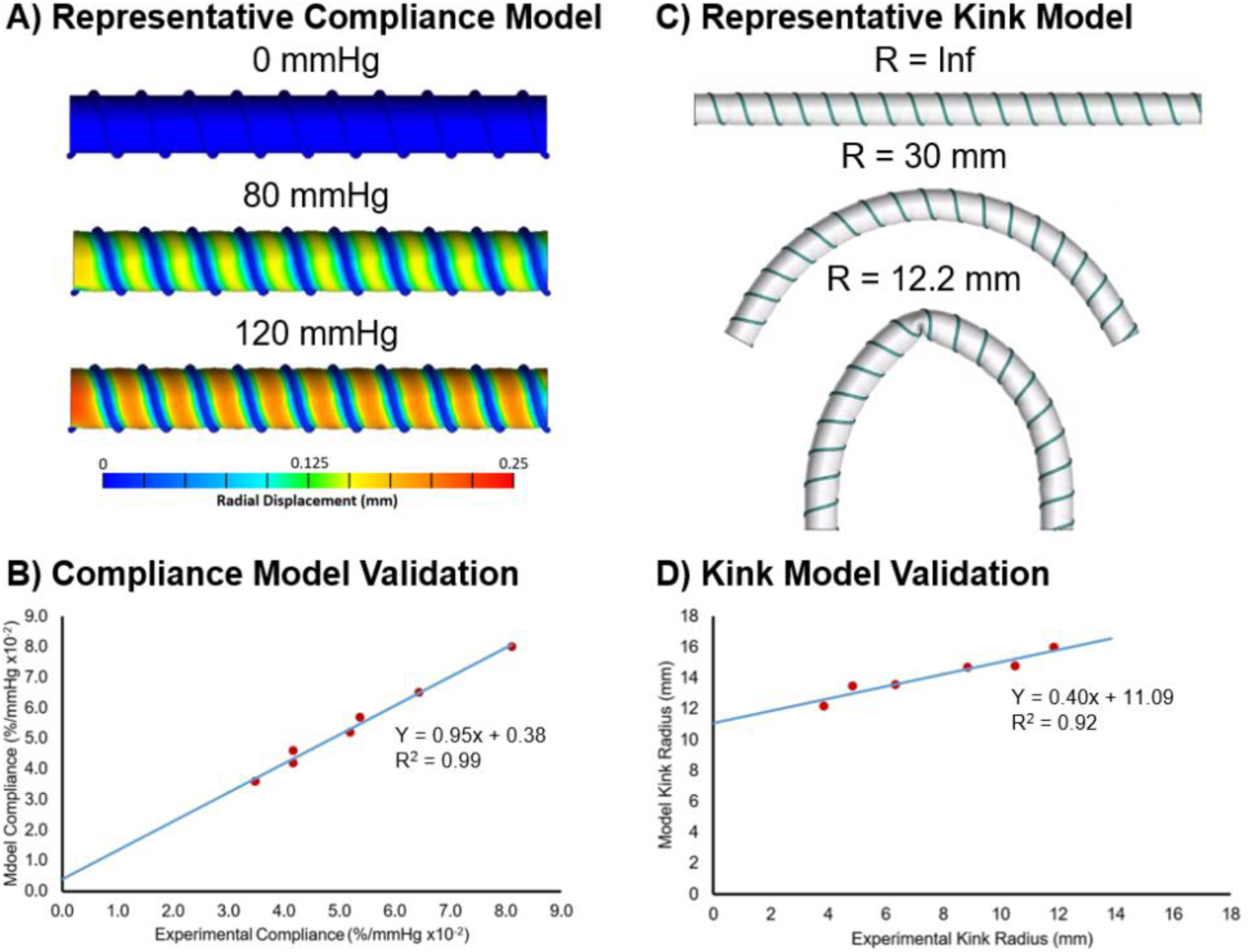
Results of the computational simulations for graft compliance and kink radius. A) Representative compliance model simulation at pressures of 0 mmHg, 80 mmHg, and 120 mmHg. B) Linear regression for validation of compliance model. C) Representative graft kink radius model at T = 0, 1.5, and 2.6 showing the bending progression of the graft and kink formation. D) Linear regression for validation of kink radius model.

The validated computational simulations allow for parsing of the design space to identify graft designs that balance graft compliance and kink resistance. Although the computational simulations have the ability to explore the complete design space of the grafts, structure-property relationships and compliance simulations performed previously revealed that a large portion of the design space does not increase graft compliance beyond previously identified graft.^22^ As an initial assessment, a reduced design space of 11 candidates was selected for *in silico* screening. The reduced design space consisted of coil spacings of 1.0 - 2.0 mm, coil diameters of 0.15-0.30 mm, and a coil modulus of 2.9 MPa. Tighter coil spacings were identified from testing as necessary to maintain adequate kink radius. Thus, a coil spacing below 2.5 mm was chosen in 0.5 mm increments to include 1.0 mm, 1.5 mm, and 2.0 mm. The compliance model indicated that coil diameters of 0.30 mm and below can achieve compliances of at least 5.0 %/mmHg x 10^-2^. Although possible to fabricate a coil with a diameter of 0.10 mm, structure-property relationships indicated that this design was unlikely to achieve the target kink radius, and thus diameters ranging from 0.15 mm to 0.30 mm in diameter in 0.05 mm increments were chosen. A coil modulus of 2.9 MPa was selected as it provided higher compliances as compared to the 8.1 MPa coil. Grafts that did not meet target compliance based on the preliminary compliance simulation were eliminated, resulting in 11 design combinations, **Table 1**. The FE model predictions indicated that three of these graft designs have a predicted compliance greater than 5.0 %/mmHgx10^-2^, but none of the designs simulated had improved kink resistance as compared to the previously identified graft, **Table 1**. This was confirmed with fabrication and kink radius testing of a candidate graft.

**Table 1:** Model-directed design space and resulting computational simulation results.

| Graft Design | Modulus (MPa) | Spacing (mm) | Diameter (mm) | Model Compliance (%/mmHg x 10<sup>-2</sup>) | Model Kink Radius (mm) |
| --- | --- | --- | --- | --- | --- |
| 1 | 2.9 | 1.0 | 0.15 | 4.9 | 14.9 |
| 2 | 2.9 | 1.0 | 0.20 | 4.0 | 14.7 |
| 3 | 2.9 | 1.0 | 0.25 | 3.2 | 15.0 |
| 4 | 2.9 | 1.5 | 0.15 | 5.7 | 14.5 |
| 5 | 2.9 | 1.5 | 0.20 | 4.8 | 14.3 |
| 6 | 2.9 | 1.5 | 0.25 | 4.1 | 14.6 |
| 7 | 2.9 | 1.5 | 0.30 | 3.6 | 14.5 |
| 8 | 2.9 | 2.0 | 0.15 | 6.2 | 15.1 |
| 9 | 2.9 | 2.0 | 0.20 | 5.5 | 14.9 |
| 10 | 2.9 | 2.0 | 0.25 | 4.8 | 15.5 |
| 11 | 2.9 | 2.0 | 0.30 | 4.3 | 15.3 |

## 4.0 Discussion

Despite significant efforts, synthetic vascular grafts have yet to achieve long-term success in small diameter applications such as coronary artery bypass grafting. An estimated 40% of small diameter graft failures within 5-years have been attributed to intimal hyperplasia and is hypothesized to develop as a result of compliance mismatch between the graft and native vasculature.^2^ We previously utilized iterative design of an electrospun polyurethane to achieve improved compliance matching while maintaining requisite burst pressure and suture retention strength for clinical use. Although this high compliance graft demonstrated promising results in the prevention of intimal hyperplasia in an *ex vivo* models, the graft failed in large animal testing due to kinking.^3^ Herein, we developed advanced manufacturing to generate a coil-supported multilayer graft to impart improved kink resistance.

A matrix of coil-supported multilayer grafts were fabricated with discrete coil spacings, diameters, and moduli and tested to establish structure-property relationships between coil design and graft compliance and kink resistance. There was generally an inverse relationship between kink resistance and compliance in the coil-supported grafts with parameters that increased kink resistance also resulting in a reduction in compliance. The decrease in compliance with tighter coil spacing was attributed to the increase support points of the coil progressively restricting the graft circumferential expansion. Similar results were reported by Adhikari et al. where a tighter polyethylene terephthalate coil increased the graft circumferential elastic modulus.^2,35^ They attributed this to the circumferential modulus being increasingly dictated by the coil mechanical properties at tighter spacings instead of the electrospun mesh.^35^ The reduction in kink radius with tighter coils was hypothesized to be due to the increased number of support points along the graft length. Wu et al. also reported a significant improvement in kink resistance in a tighter-spaced coil.^36^ In contrast to the compliance decrease that was uniform over the spacings tested, the kink resistance sharply improved at 2.5 mm spacing compared to the 5.0 and 7.5 mm spacings, demonstrating a non-linear relationship between the number of support points and the kink resistance.

The reduction in graft compliance and improved kink resistance with increased coil diameter was attributed to the increased cross-sectional area of the coil reducing graft circumferential expansion. In contrast to coil spacing, the effect of coil diameter on both compliance and kink radius was more linear across the range investigated. Interestingly, other researchers have seen minimal trends in kink resistance with regards to coil diameter.^15^ Wu et al., reported minimal changes to graft kink radius when changing the supporting poly-ε-caprolactone (PCL) coil diameter.^15^ This difference may be attributed to innate polymer differences such as the elastic and bending modulus of the supporting coil with PCL typically being greater than that of the polyurethane coil utilized here.^27,37-40^ Thus, even a smaller diameter PCL coil may be able to sufficiently support the graft during bending as compared to a polyurethane coil. Indeed, minimal effects of coil modulus were observed for the range tested here.

The establishment of structure-property relationships enabled the identification of a graft design that was within commercial graft kink radii (graft =4.8 ± 1.5 mm, clinical grafts ≤ 5.6 mm) and had a compliance comparable to the saphenous vein (graft = 4.1 ± 0.4%/mmHg x 10^-2^, saphenous vein = 4.4 %/mmHg x 10^-2^).^31,32^ This combination of properties is promising toward developing an off-the-shelf graft that may provide improved treatment options for the 20% of patients annually that undergo multiple bypass or that do not have a suitable peripheral vessel.^3,41^ Although this graft has potential to expand treatment options for patients, it is anticipated that higher compliances will further improve long-term outcomes. For example, prior work by Post et al. explored the impacts of synthetic graft compliance on the development of intimal hyperplasia in an *ex vivo* porcine carotid artery model. The work demonstrated that grafts of increasing compliance reduced early markers of intimal hyperplasia.^3^ Although improved compared to the ePTFE representative graft, a graft that had a compliance comparable to the saphenous vein still displayed early markers of intimal hyperplasia.^3^ Clinical studies further corroborate this result with saphenous vein grafts (compliance = 4.4 %/mmHg x 10^-2^) having failure rate of 45% after 5-12 years compared to the higher compliance internal mammary artery (compliance = 11.5 %/mmHg x 10^-2^) with 97% of grafts remaining patent in the same timeframe.^30,42,43^ A graft with a higher compliance than the saphenous vein and a kink radius within current commercial graft ranges is hypothesized to further enhance long-term patient outcomes. However, iterative approaches to achieve these criteria are time prohibitive due to the extensive design space.

To accelerate the design of the coil-supported graft, an FE framework to predict graft computational simulations for kink radius and compliance was previously developed by Jiang et al.^22^ In combination, these simulations allow for a rapid probing of the design space to identify an optimized design that surpasses the compliance of the previously identified graft while maintaining a kink resistance within commercial graft ranges. The graft compliance model agreed well with the experimental results (<11% difference). A linear regression further demonstrated the agreement as the slope was close to 1.0 (slope = 0.95) and had an R^2^ value of 0.99, demonstrating a close fit for the trend in the results. This high level of agreement allows for the model to be used to accurately predict the numerical value of the graft compliance for design optimization. The graft kink model overpredicted the numerical value of graft kink radius. For example, a graft with a kink radius of 3.8 mm experimentally had a kink radius of 12.2 mm for the model. Although the model could predict the trends of the impact of coil design on kink radius, the slope of 0.40 demonstrated that it also underpredicted the extent of the effect of coil parameters on kink radius. Based on this additional validation of the model values with experimental graft results, we expected that the models provided sufficient predictive capability for *in silico* optimization.

Previous evaluation of 100 graft designs using a validated FE model to quantify graft compliance, kink radius, and buckling load indicated that a portion of the design space was unlikely to achieve a target kink radius and superior compliance to the previously identified graft.^22^ Due to this, a reduced design space with the highest potential of achieving a superior compliance and maintaining a suitable kink radius was explored. The model predicted that three of the 11 candidate designs had a compliance above 5.0 %/mmHg x 10^-2^; however, none of the designs simulated had improved kink resistance as compared to the previously identified graft. Although this result did not yield an optimized graft design to date, this finding is significant as it saved substantial experimental time in fabrication and graft testing and validates the computational framework for accelerating graft design. By exploring a larger design space and additional coil geometries, full optimization of the graft could be performed to achieve a graft with arterial compliance matching while maintaining kink resistance and high burst pressure. To this end, improving the FE model to predict kink radius could promote agreement between the experimental and simulation results. As the developed computational framework is modular and scalable, integrating model advances into the batch processing scheme is feasible. As examples, more advanced material models for the electrospun polyurethane graft could be incorporated to capture the mesh phase of the graft architecture and its anisotropic response to loading and integrating modern and certifiable uncertainty quantification techniques to capture fabrication variability (e.g., material imperfections) would promote FE model robustness and confidence in model predictions.^8,22^ In addition, incorporating a non-uniform sampling across the parameter design space (e.g., weighted approximate Fekete points) could better capture promising parameter spaces and promote optimization efficiency.^44^ Of course, these additions would inevitably raise the computational expense. Yet, they are necessary to advance the framework’s functionality and ability to promote validation of the FE model and better differentiate the effect of coil design on kink radius.

## 5.0 Conclusion

In this work, we developed a coil-reinforced multilayer vascular graft design to achieve a balance of high compliance and kink resistance. Due to the inverse relationship between compliance and kink resistance, structure-property relationships of coil design were first established. Closer coil spacings achieved desired kink resistance but had a large negative impact on graft compliance. Smaller diameter coils were able to recapture some of the reduced compliance but at the consequence of reduced kink resistance. In contrast, the range of coil modulus tested had relatively minimal effect on graft properties. Although each parameter had an inverse effect on compliance and kink resistance, the effects were of different magnitude which indicates that potential to optimize parameter combinations to achieve target biomechanical properties. A candidate graft was identified with a compliance, burst pressure and suture retention strength that meets or exceeds the saphenous vein and a kink radius comparable to current clinical grafts. To further improve graft compliance, previously developed FE models were leveraged to parse through a larger design space and accelerate graft design and testing. The developed compliance and kink radius models were validated with the additional graft experimental results from this study indicating good prediction of graft biomechanical properties. Although this initial parameter range tested did not yield a graft that improved upon the previous balance of graft properties, this combination of advanced manufacturing and computational framework paves the way for future model-driven design to further optimize graft performance.

. In addition to providing an improved synthetic graft for small diameter applications, these studies establish the potential of this model-directed approach to accelerate vascular graft design.

## Acknowledgment

This work was supported by the American Heart Institute Predoctoral Fellowship (23PRE1020036—https://doi.org/10.58275/AHA.23PRE1020036.pc.gr.161179), NSF GRFP (2023355666), and the National Institutes of Health (R01 HL150608). Bionate® 80A and 55D were provided by DSM Biomedical (Exton, PA).

## Supplemental Information

**Figure S1:**
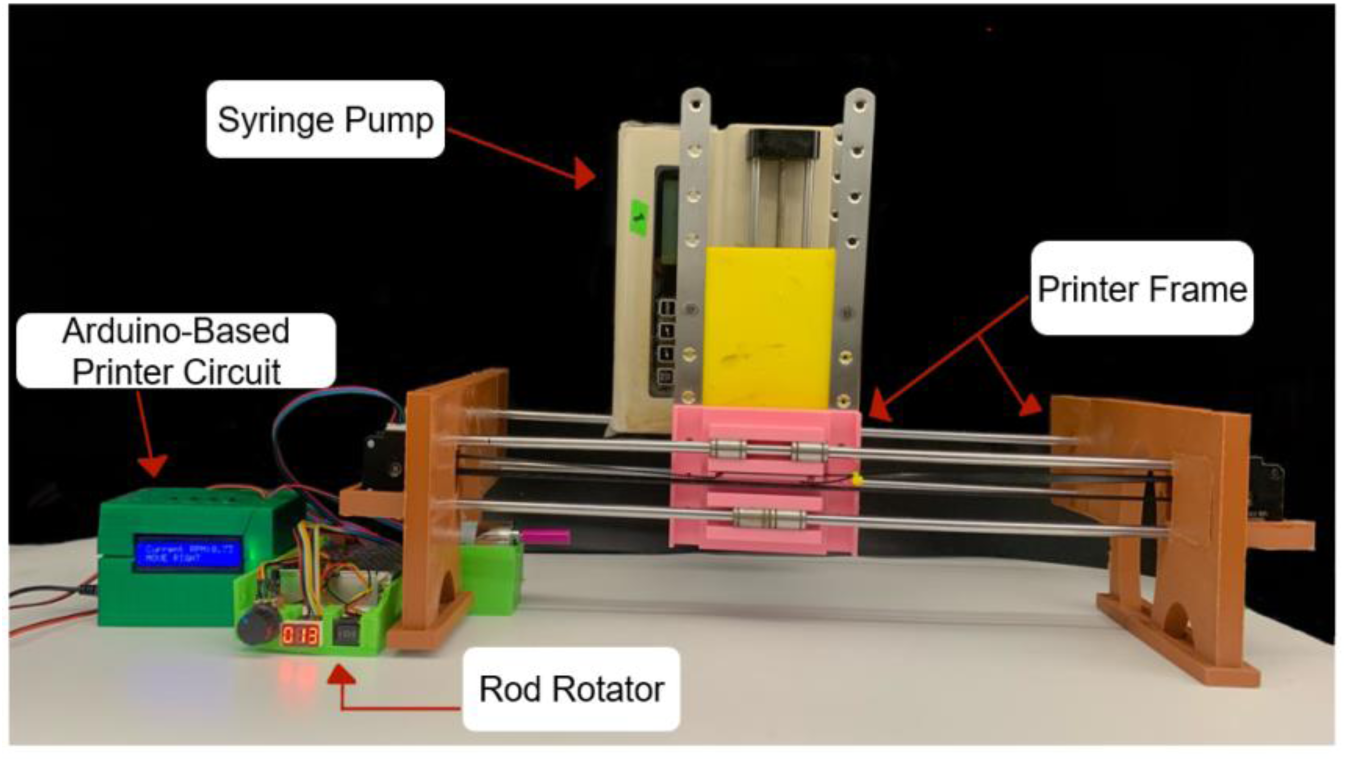
Custom coiling printer. A 3D printed frame held a syringe pump with lateral motion controlled via a stepper motor and Arduino.

**Figure S2:**
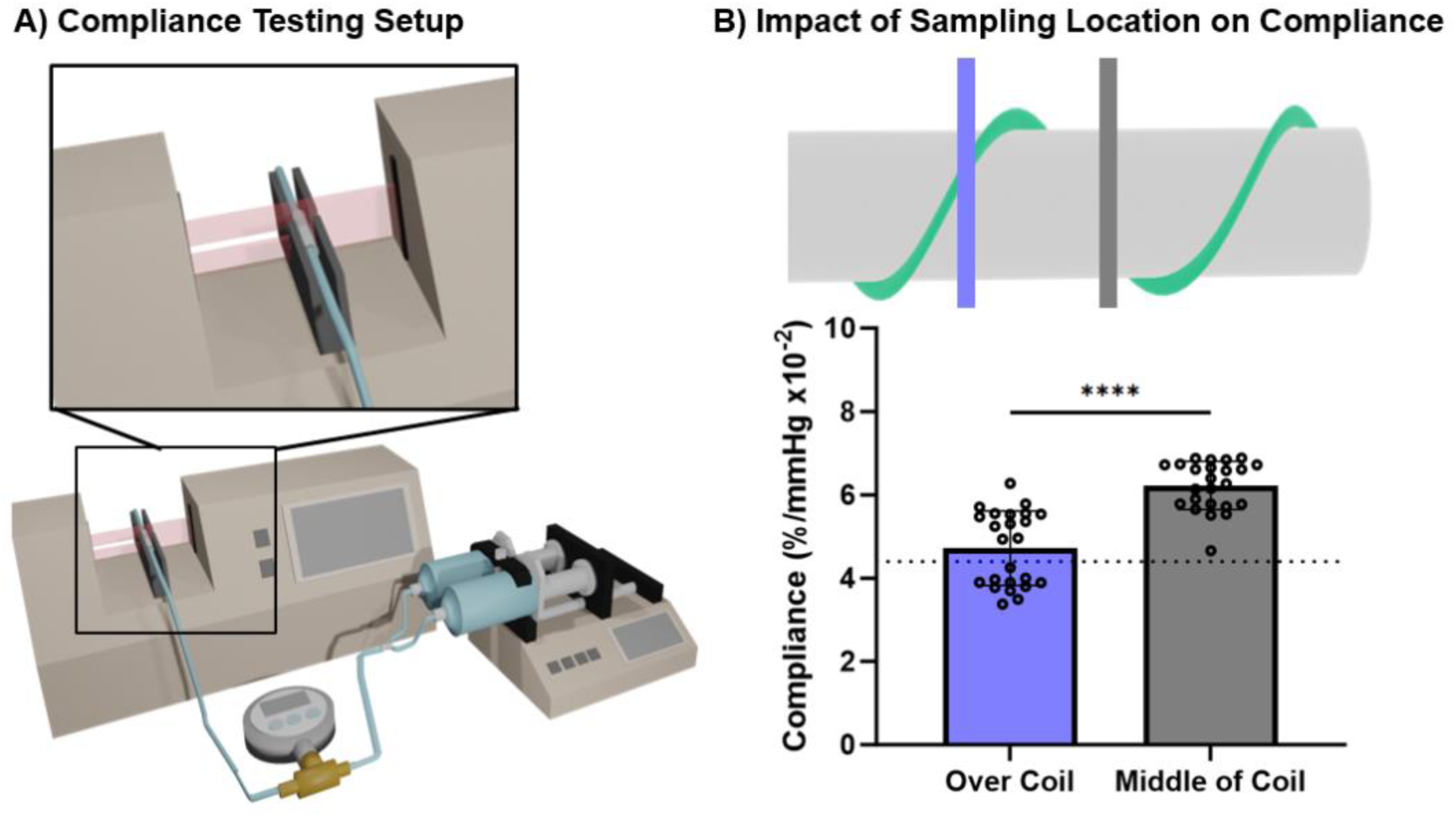
Effect of graft positioning and compliance. A) Depiction of compliance measurement testing setup. B) Sampling location and the effect on compliance.

**Table S1:** Printer settings for target graft designs.

| **Coil**<br>**Polymer/**<br>**modulus** | **Coil**<br>**Spacing**<br>**(mm)** | **Coil**<br>**Diameter**<br>**(mm)** | **Lateral**<br>**Speed**<br>**(mm/min)** | **Graft**<br>**Rotational**<br>**Speed (RPM)** | **Needle**<br>**Gauge** | **Flow**<br>**Rate**<br>**(ml/hr)** |
| --- | --- | --- | --- | --- | --- | --- |
| Bionate® 55D<br>8.1 MPa | 2.5 | 0.35 | 29 | 10 | 18 | 10 |
| Bionate® 55D<br>8.1 MPa | 5.0 | 0.35 | 49 | 10 | 18 | 10 |
| Bionate® 55D<br>8.1 MPa | 7.5 | 0.35 | 82 | 10 | 18 | 10 |
| Bionate® 55D<br>8.1 MPa | 2.5 | 0.25 | 29 | 10 | 18 | 5.4 |
| Bionate® 55D<br>8.1 MPa | 2.5 | 0.15 | 29 | 10 | 20 | 30 |
| Bionate® 80A<br>2.9 MPa | 2.5 | 0.35 | 29 | 10 | 16 - 18 | 10 |

